# Efficient Multiplex Genome Editing Tools identified by Protoplast Technology in *Phalaenopsis*

**DOI:** 10.1101/2020.09.29.315200

**Authors:** Keke Xia, Dengwei Zhang, Guangyu Liu, Xiaojing Xu, Yong Yang, Guo-Qiang Zhang, Hai-Xi Sun, Ying Gu

**Affiliations:** BGI-Shenzhen, Shenzhen 518083, China; Guangdong Provincial Key Laboratory of Genome Read and Write, BGI-Shenzhen, Shenzhen, 518120, China; Laboratory for Orchid Conservation and Utilization, The Orchid Conservation and Research Center of Shenzhen, The National Orchid Conservation Center of China, Shenzhen 518114, China

**Keywords:** *Phalaenopsis*, protoplast, Cas9, Cpf1, orchid, multiplex genome editing

## Abstract

*Phalaenopsis* orchids are popular ornamental plants worldwide. The application of the efficient multiplex genome editing tools in *Phalaenopsis*, will greatly accelerate the development of orchid gene function and breeding research. In this study, we establish a fast and convenient *Phalaenopsis* protoplast platform for the identification of functional genome editing tools. Two multiplex genome editing tools, PTG-Cas9 (PTG, polycistronic tRNA gRNA) system and PTGm-Cas9 (PTG-Cas9 system with modified sgRNA structure) system are designed to edit *PDS* gene of commercial *Phalaenopsis* ST166 at four target sites. We find that both PTG-Cas9 and PTGm-Cas9 system are functional in *Phalaenopsis*, and the PTGm-Cas9 system with modified sgRNA has a higher editing efficiency than PTG-Cas9 system. Further, we design another multiplex genome editing tool, termed as DPII-Cpf1 system (dual Pol II promoter to drive the expression of Cpf1 endonuclease and crRNA), to edit *PDS* gene of *Phalaenopsis* at four target sites likewise. All the four targets are efficiently edited by DPII-Cpf1 system, and the total mutation rate is about 3 times higher than that of PTGm-Cas9 system. Taken together, using the *Phalaenopsis* protoplast platform, we successfully establish two efficient multiplex genome editing tools for *Phalaenopsis* research, PTGm-Cas9 and DPII-Cpf1. The multiplex genome editing tools established in this study have great application potentials in efficiently constructing large-scale knockout mutant libraries of orchid and speeding up orchid precise breeding.

## Introduction

*Phalaenopsis* species are worldwide popular ornamental plants and are of enormous value to commercial horticulture and plant scientific research^1^. However, due to the long growth cycle and the lack of efficient genetic transformation and genome editing technologies on Orchidaceae plants, the investigation into orchid genome function and the breeding of new orchid species are seriously hindered. In recent years, the fast progresses in genomics, including sequencing technology and genome editing technology such as CRISPR/Cas (Clustered Regularly Interspaced Short Palindromic Repeats/CRISPR-associated endonuclease) system, provide valuable genome sequences of Orchidaceae plants and powerful tools for rapid orchid breeding and orchid horticulture research.

For the past few years, the draft genome sequences of two Orchidacease, *Phalaenopsis equestris*^1^ and *Dendrobium catenatum* Lindl.^2^, are released. And a large number of functional gene families of orchid have been excavated, such as crassulacean acid metabolism (CAM) genes, MADS-box genes, disease resistance genes and heat-shock protein genes. These findings provide key resources for further studies on orchid gene function and orchid genetic improvement. Editing and obtaining the mutants of these functional genes is one of the necessary approaches for further orchid researches.

The CRISPR/Cas genome editing technology is a simple and efficient system for gene knockout and other genetic manipulation, and it has been successfully applied to various plant species^3–9^, including *Phalaenopsis equestris*^10^ and *Dendrobium officinale*^11^. The diverse and expanding CRISPR toolkits are flexible and efficient to achieve precise genome editing and transcriptome regulation^12,13^. Moreover, researchers could obtain transgene-free mutant plants based on the CRISPR/Cas system^14–16^. Particularly, many versatile and multiplexed CRISPR expression systems have been developed, such as single transcript unit system^17,18^, polycistronic tRNA– gRNA system^19,20^, and HH-gRNA-HDV system^20–22^, enabling to edit multiple genes simultaneously without sacrificing the length of expression cassettes and the editing efficiency. The flexible multiplex CRISPR toolkits are ideal approaches to efficiently and high-throughput obtain multiple mutants of orchid for breeding and horticulture research. However, up to now, there is only one study that successfully generated mutants of *MADS* genes in *Phalaenopsis equestris* using CRISPR/Cas9 system^10^. In contrast, no report has been published for the use of CRISPR/Cpf1 system in orchid research, largely due to the difficult genetic transformation in *Phalaenopsis*. Therefore, the establishment of a platform in Orchidaceae to rapidly screen functional and efficient CRISPR/Cas toolkits is urgently needed.

Plant protoplasts-based platform is a reliable and convenient strategy for plant science researches^23^. Transient protoplast transfection technology has been widely used to investigate gene regulations, and to study the subcellular localization and interaction of proteins, and is also an alternative strategy to screen efficient CRISPR toolkits in plants^24–26^. Thus far, orchid protoplasts have been successfully isolated from *Dendrobium*^27–29^, and *Phalaenopsis*^30–32^, and the orchid protoplast transient expression system has been established in *Phalaenopsis aphrodite subsp. formosana* (m1663)^31^, *Phalaenopsis* hybrid cultivar ‘Ruili Beauty’^32^, and *Cymbidium* orchid^33^. However, these orchid protoplast transient expression technologies have not been applied to screen efficient CRISPR-Cas toolkits for orchid research.

In this study, we established a seedling-leaf protoplast-based platform to rapidly identify functional and efficient multiplex genome editing systems in *Phalaenopsis*. We found that the PTG-Cas9 multiplex genome editing tool was effective in *Phalaenopsis*, and replacing the classical sgRNA scaffold with a modified sgRNA scaffold, could further increase the editing efficiency. In addition, we built an efficient DPII-Cpf1 system for *Phalaenopsis* multiplex genome editing. The efficient toolkits developed in this study would facilitate the construction of large-scale mutant libraries and therefore promote the development of precise breeding of orchids.

## Results

### Assessment of PTG-Cas9 multiplex genome editing system in *Phalaenopsis* via protoplast technology

CRISPR/Cas genome editing technologies, especially the multiplex genome editing toolkits, have great potential to be used to build large-scale knockout mutant library and to investigate gene function and then to facilitate breeding research. However, due to a deficiency of the rapid screening technology for orchid, researchers are unable to optimize the gene editing tools for Orchidaceae plants. To overcome this obstacle, we established a convenient and efficient transient protoplast platform in commercial *Phalaenopsis* ST166. Two-month-old seedling leaves of aseptic *Phalaenopsis* ST166 leaves, cultured in multiplication medium, were cut into 0.5~1.0-mm strips and digested by protoplast isolation solution (PIS). After only 3 hours of enzymolysis, protoplasts were washed and prepared for transfection (Fig. 1). To estimate the transfection efficiency, the plasmid pS1300-GFP (Fig. S1a) that expresses green fluorescent protein (GFP) was transfected into protoplast via PEG mediated plasmid transformation method. The GFP signals were detected in about 55% of protoplasts (Fig. S1b, c). The *Phalaenopsis* protoplast transient expression technology was then used for the following gene editing experiments.

**Fig. 1.**
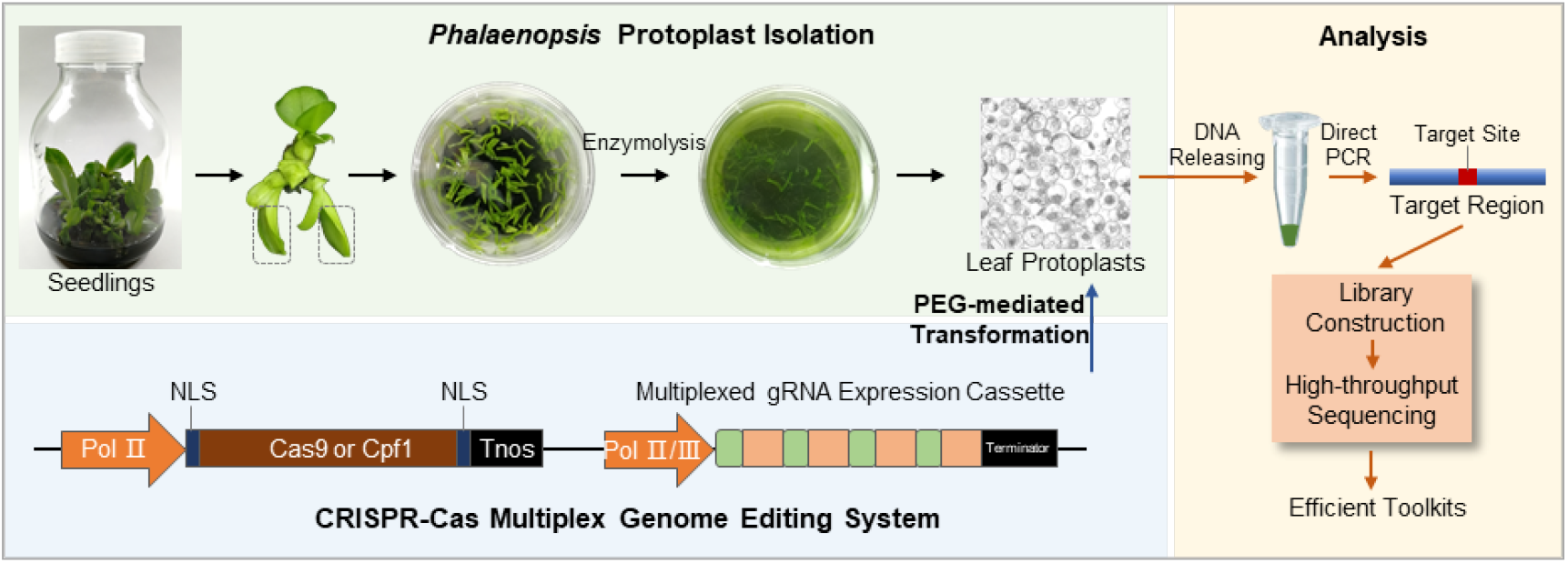
Schematic diagram of efficient multiplex genome editing toolkits screening in *Phalaenopsis* protoplast. The CRISPR-Cas based multiplex genome editing plasmids were delivered into protoplasts isolated from proliferation cultured seedling leaves of *Phalaenopsis*, by PEG-mediated protoplast transfection. After 48 h of incubation, the protoplasts were collected and the genome DNA was released for direct PCR amplification. The target regions were purified for library construction and high-throughput sequencing, and the data was used for the editing efficiency analysis.

tRNA-based multiplex genome editing tools have been wildly used in plant researches, but not in Orchidaceae. To assess the feasibility of these tools in orchid plants, we cloned a fragment of *PDS* gene of *Phalaenopsis* ST166, according to the published genome sequences of *Phalaenopsis equestris*, and designed PTG-Cas9 multiplex genome editing system that contains four gRNAs targeting the *PDS* gene. The transcription of gRNA cassette was driven by OsU3 promoter (Fig. 2a). Target sites of *PDS* were shown in Fig. 2b. The designed PTG-Cas9 system was delivered into protoplasts of ST166 to examine the editing effect. Two days after transfection, the genomic DNA of protoplasts was released, and the target regions were PCR-amplified, followed by library construction and high-throughput sequencing (Fig.1a). For library construction, the target region of 1&2 and 3&4 were amplified separately, using primers F1 and R1, and F2 and R2, respectively (Fig. 2b). We successfully detected editing events in the sequencing data. As shown in Fig. 2c, representative reads carrying mutations of target 1, 2, 3 and 4 were listed, and insertion, deletion, and substitution editing events were detected at all of the four target sites. These results showed that the designed PTG-Cas9 multiplex genome editing system was functional and all the designed target sites could be successfully edited in *Phalaenopsis*.

**Fig. 2.**
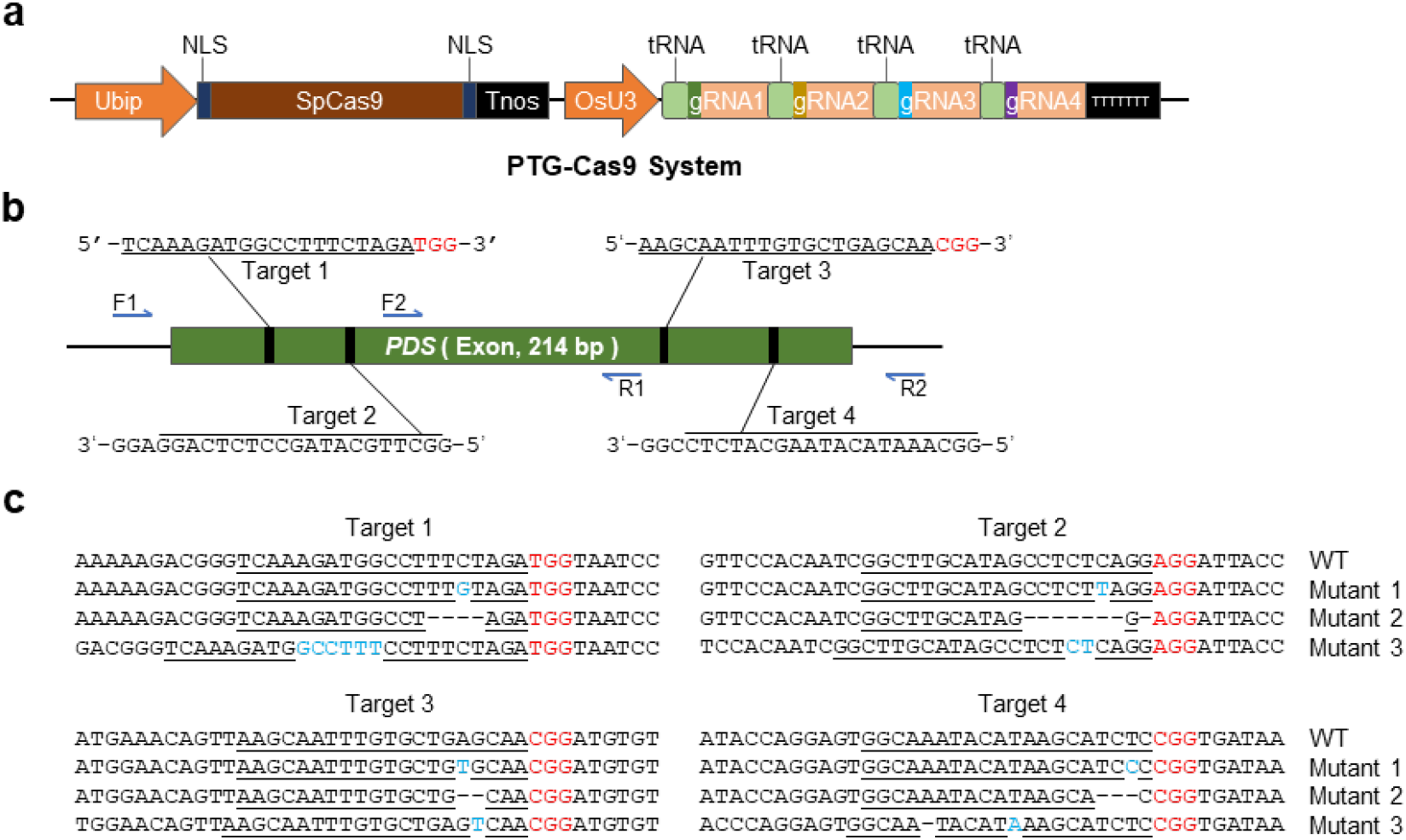
PTG-Cas9 multiplex genome editing system is effective in *Phalaenopsis*. **a** The architecture of PTG-Cas9 system. The tRNA-mediated multi-gRNAs expression cassette is driven by OsU3 promoter. **b** The illustration of the four targeted sites of *PDS* gene edited by PTG-Cas9 system. The primers used for target region amplification is indicated with blue arrows. F1 and R1 primers were used for the amplification of targets 1 and 2. F2 and R2 primers were used for the amplification of targets 3 and 4. **c** The mutations at four target sites edited by PTG-Cas9 system. The plasmid showed in **a** was delivered into *Phalaenopsis* protoplasts, and the editing results were analyzed by high-throughput sequencing. Mutations were listed as representatives. The PAM sequences are highlighted in red. The target sequences are marked by underlines. The insertion or mutation bases are shown in light blue.

### Improvement of PTG-Cas9 editing efficiency with modified sgRNA structure

Single-guide RNA (sgRNA) is indispensable for CRISPR/Cas9 system. Therefore, optimizing sgRNA structure is a feasible way to improve the efficiency of CRISPR-Cas9 system^34–36^. Here, to improve the efficiency of PTG-Cas9 system, the classical sgRNA scaffold was swapped for a modified sgRNA scaffold, that has been shown to improve editing efficiency in TZM-bl cells and rice^35,36^, and the modified PTG-Cas9 system was termed as PTGm-Cas9 system. As shown in Fig. 3a, the original sgRNA was modified by extending the duplex and mutating continuous sequence of Ts, a potential transcription pause site, at position 4 to C. To investigate the editing efficiency of PTGm-Cas9 system, plasmids expressing PTGm-Cas9 system or PTG-Cas9 system were transformed into protoplasts of *Phalaenopsis* ST166 separately, and the target regions were amplified for the library construction and sequencing. The analysis results showed that, in T1 (one of the three independent experiments), the mutation rates of the four target sites were 1.15%, 0.71%, 0.74%, 1.29% respectively, totaling 3.89%, when using PTG-Cas9 system. In contrast, the corresponding mutation rates were 1.44 %, 0.82%, 0.83%, 1.38% respectively, totaling 4.47%, when using PTGm-Cas9 system (Fig. 3b). The editing efficiency was improved in PTGm-Cas9 system compared to that in PTG-Cas9 system. And this observation was reproducible when the other two batches were tested (Fig. 3b, S2). Overall, our data suggested that the modified sgRNA structure was also capable of improving the editing efficiency of CRISPR/Cas9 system in *Phalaenopsis*, and the tRNA-based multiplex editing tools for *Phalaenopsis* could also be improved by optimizing the sgRNA structure.

**Fig. 3.**
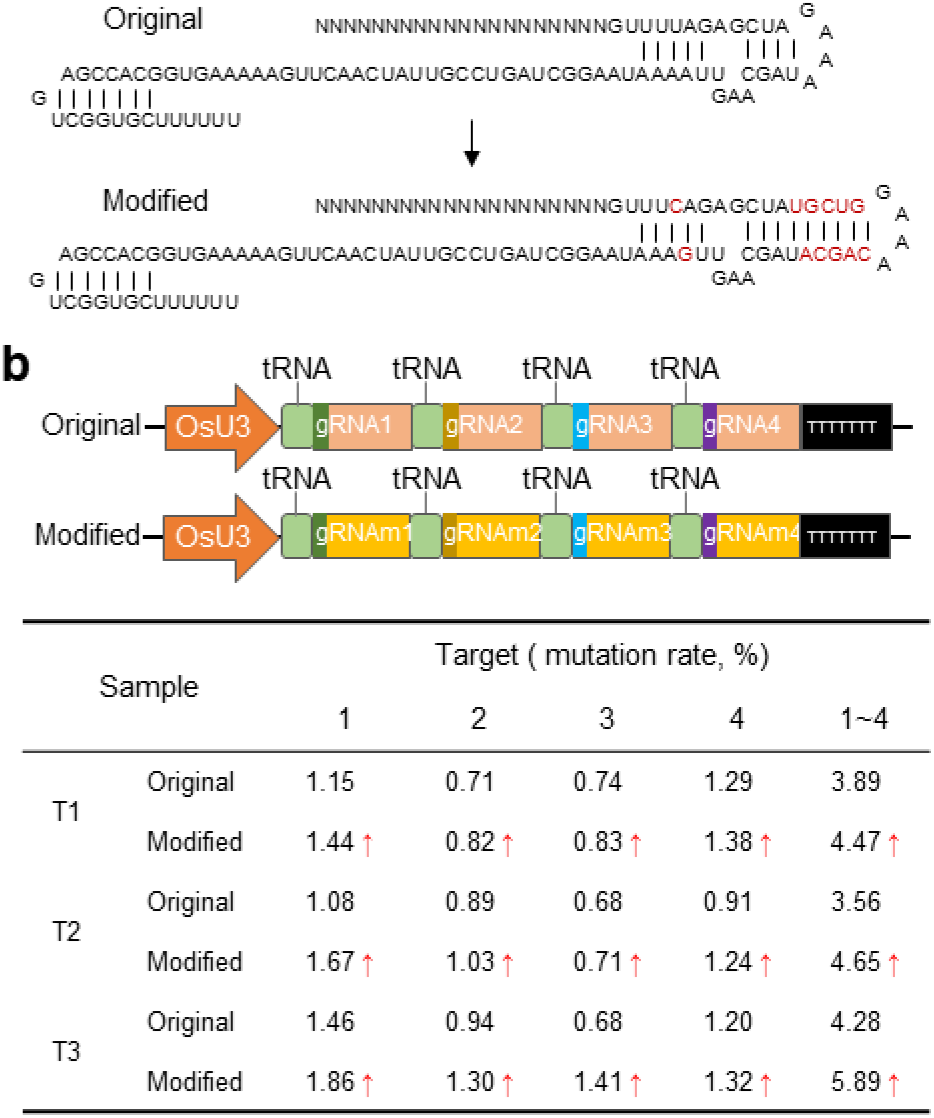
PTGm-Cas9 system improves the editing efficiency in *Phalaenopsis*. **a** Schematic representation of original and modified sgRNA structure. The duplex extension and mutation in modified sgRNA are highlighted in red. **b** The mutation rate of four target sites of *PDS* gene edited by PTG-Cas9 or PTGm-Cas9. The architecture of multi-gRNAs expression cassette in PTG-Cas9 or PTGm-Cas9. The original and modified sgRNA are shown in orange and yellow respectively. The data in table showed the mutation rate of the four target sites respectively and summarized. T1, T2, and T3 indicate three independent experiments. The arrows indicate the improved mutation rate. The sequencing of the amplicons was repeated 3 times, using genomic DNA from three independent protoplast samples. Mutation rate was calculated as the ratio of the number of mutant reads to that of the total reads.

### Assessment of DPII-Cpf1 multiplex genome editing system in *Phalaenopsis* via protoplast technology

Compared to CRISPR/Cas9 system, Cpf1 endonuclease has a smaller molecular weight, and requires shorter CRISPR RNA (crRNA)^37,38^. These advantages of CRISPR/Cpf1 system could help to reduce the overall size of the plant transformation vector, making it more suitable for multiplexed genome editing for plants^39^. However, the CRISPR/Cpf1 system has not been applied in Orchidaceae so far. Here, we developed a multiplex genome editing tool for *Phalaenopsis*, named DPII-Cpf1 (dual Pol II promoter-Cpf1) (Fig. 4a). The *Phalaenopsis* codon-optimized LbCpf1 is driven by Super promoter, and the multi-crRNA expression cassette contains a double ribozyme^21^ as well as four DR-guide units^40^ that each contains 21 bp of DR sequence and 23 bp of guide sequence. *Cestrum Yellow Leaf Curling Virus* (CmYLCV)^41^ promoter and Poly (A) signal were utilized for multi-crRNA expression and transcription termination respectively. Similarly, four target sites of *PDS* gene were selected for simultaneous targeting by the designed DPII-Cpf1 system (Fig. 4b). The four target regions were respectively amplified from the genomic DNA of protoplasts with or without transfection, with primers F3 and R3. The sequencing results (Fig. 4c) showed that the mutation rates of four target sites were 1.04%, 3.80%, 4.13%, and 9.63% respectively, totaling 18.60%, in T1. The editing events were detected at all of the four target sites. And this observation was further confirmed by other two batches of testing, T2 (2.79%, 1.23%, 1.84%, 7.87%, totaling 13.73%) and T3 (3.29%, 1.39%, 1.84%, and1.93%, totaling 14.44%). Our data indicated that DPII-Cpf1 multiplex genome editing system was also functional and all the designed target sites could be successfully edited in *Phalaenopsis*.

**Fig. 4.**
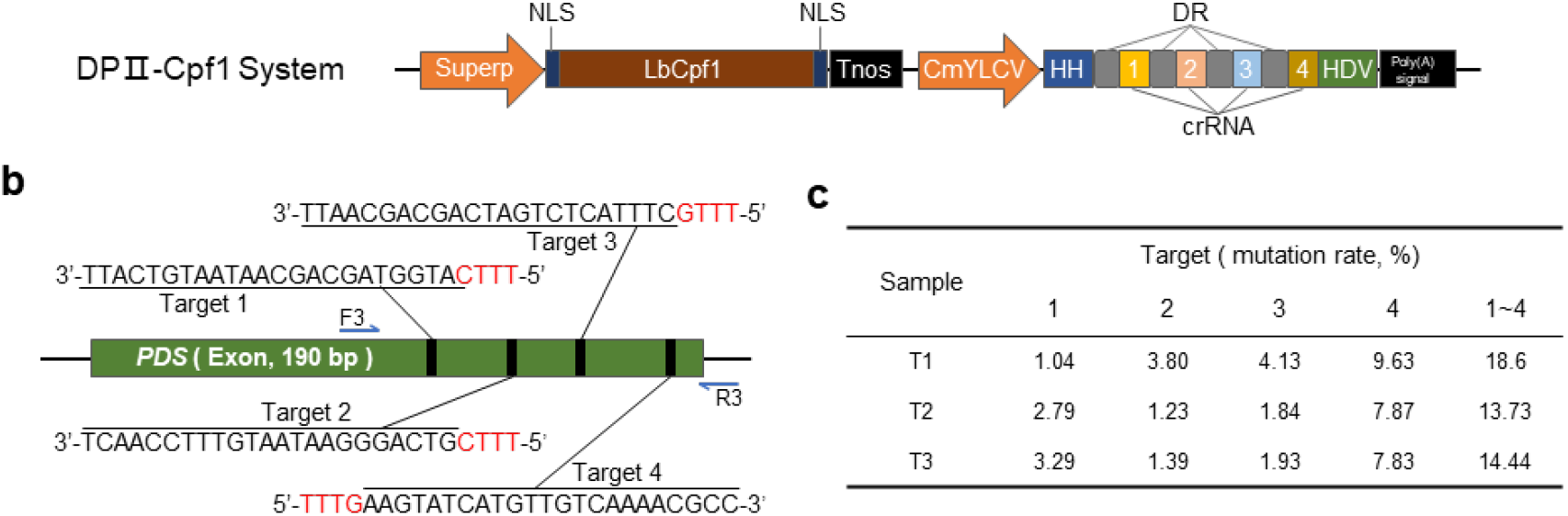
DPII-Cpf1 multiplex genome editing system is effective in *Phalaenopsis*. **a** The architecture of DPII-Cpf1 system. The ribozyme-based multi-crRNA expression cassette was driven by CmYLCV promoter. HH, hammerhead ribozyme; HDV, hepatitis delta virus ribozyme; DR, direct repeat. **b** The illustration of the four targeted sites of *PDS* gene edited by DPII-Cpf1 system. The primers used for target region amplification are indicated with blue arrows. The PAM sequences are highlighted in red. The target sequences are marked by underlines. **c** The mutation rate of the four target sites edited by DPII-Cpf1 system. The plasmid showed in **a** was delivered into *Phalaenopsis* protoplasts, and the editing results were analyzed by high-throughput sequencing. Mutation rate was calculated as the ratio of the number of mutant reads to that of the total reads. T1, T2, and T3 indicate three independent experiments.

## Discussion

With the increasing knowledge on genome sequences of Orchidacease, feasible genome editing tools for these valuable plant species are urgently needed. The protoplast technology and multiplex genome editing toolkits established in this study not only provide useful tools for targeted editing in Phalaenopsis, but also hold great potentials to extend to other Orchid plants, therefore facilitating the research of orchid biology.

Transient protoplast transfection technology has been widely applied for plant biology researches. Orchid protoplast had been successfully isolated from *Dendrobium*^27–29^, and *Phalaenopsis*^30–32^, using the suspension cells, leaves, flower petals, or callus. Until recently, the orchid protoplast transient expression system was reported only in two *Phalaenopsis* species, *Phalaenopsis aphrodite subsp. formosana* (m1663)^31^ and *Phalaenopsis* hybrid cultivar ‘Ruili Beauty’^32^, and Cymbidium orchid^33^. However, compared to the protoplast technology established in this study, these protoplast isolation systems are suboptimal. One requires young leaves of shoots induced from flower nodal buds, and the others require fully open flower petals. These materials are not readily available throughout the year, which would seriously impede the sustainability of orchid researches. In this study, the protoplasts were isolated from seedling leaves of *Phalaenopsis* ST166, cultured in multiplication medium, which could ensure an adequate source of experimental materials throughout the year. Moreover, in this study, the enzymolysis time is only 3 h, which is much shorter than the reported 6~16 h^31,32^. These advantages will significantly shorten the experimental period and improve the efficiency of research. In the future, with the help of a combination of high-throughput CRISPR screening technologies^42^ and protoplast transfection and regeneration technology^28,30^ in Orchidacease, researchers could extensively establish the large-scale orchid mutant library and provide abundant resources for orchid gene function and breeding studies.

In addition, the direct PCR method was developed in this study to amplify target sequences form *Phalaenopsis* protoplast without genomic DNA extraction, providing a great convenience to the efficiency analysis. Compared with previous reports that need to prepare large protoplast samples for genomic DNA extraction and PCR amplification, our protoplast direct PCR method is superior for micro samples requirements, free from DNA extraction, short experiment period, and potential to fit high-throughput screening system. This *Phalaenopsis* protoplast direct PCR method also has the potential to extend to other plant species.

tRNA processing systems are virtually conserved in all organisms. Because of the minimized sgRNA expression cassette and increased sgRNA transcript level, the editing efficiency of PTG-Cas9 system was higher than that of a single-gRNA-containing construct^26^. PTG-Cas9 multiplex genome editing tools have been applied in many plants, but not in orchid yet. Nevertheless, multiplex genome editing toolkits are more efficient, which could make up the defect of the low efficiency of stable genetic transformation in Orchidaceae. In addition, genetic redundancy is common in orchid genome. For example, MADS-box genes are important and potential gene resources for orchid flower development and modification research^43,44^. But there are 51 putative functional MADS-box genes in *Phalaenopsis equestris*, and 20 of them are highly expressed in flower tissue, while 5 are specifically expressed in flower^3^. It is a great challenge to obtain different combinations of MADS-box gene mutants simultaneously through the traditional approach to study the function of these MADS-box genes. However, the PTG-Cas9 multiplex genome editing tool, developed in this study, provides an ideal solution to solve this problem. Moreover, we also found TaU6 and OsU3 promoter showed the similar multiplex genome editing efficiency (Fig. S3), and this further enriches the PTG-Cas9 multiplex genome editing tools.

The modified sgRNA structure might enhance the ability of binding to Cas9 or increase the stability of itself to improve the editing efficiency of CRISPR/Cas9 system^35^. In this study, we found that the modified sgRNA scaffold could improve the efficiency of PTG-Cas9 multiplex genome editing system in orchid plants. However, the improvement is limited. Hu *et al.* significantly increased the editing efficiency by using modified sgRNA structure and strong endogenous promoters in rice^36^, and the efficiency of editing tools with modified sgRNA varies with different target sites^35,36^. This indicates that the PTGm-Cas9 system can be further optimized in the future, such as developing the strong orchid endogenous promoters to drive the expression of Cas9 and sgRNA cassette, using the orchid codon-optimized Cas9, and selecting more efficient target sites.

Up to now, the knowledge on the endogenous promoters of Orchidaceae is still inadequate, especially on RNA polymerase III (Pol III) promoter, such as U3 and U6 promoter. The efficiency of U3 and U6 promoter varies greatly in different plant species, and this might lead to a decrease in the efficiency of PTGm-Cas9 system. In addition, there are many limitations to Pol III‐based gRNA expression. U3 and U6 promoters are constitutive promoters, and they cannot be used to generate cell- or tissue-specific gRNA expression. In the present study, we successfully used CmYLCV, a RNA polymerase II (Pol II) promoter comparable with 35S promoter^41^, to drive the ribozyme-based multi-crRNA expression in DPII-Cpf1 system in *Phalaenopsis*. Based on DPII-Cpf1 system, the CmYLCV promoter can be replaced with orchid flower cell- or tissue-specific promoter to precisely edit the genes related to floral morphogenesis, facilitating precise breeding and floral organ development researches. In addition, in this study, DPII-Cpf1 system is about 1.4 kb less in length than that of PTGm-Cas9 system, and these features may moderate the difficulty of plasmid construction and improve the efficiency of genetic transformation. Thus, DPII-Cpf1 system could accommodate much more editing sites in one construction. Moreover, the total editing efficiency of DPII-Cpf1 system is about 4 times as much as PTGm-Cas9 system, indicating that DPII-Cpf1 system might be a much more potential multiplex genome editing tool in Orchidaceae plants. Considering that CRISPR/Cpf1 is temperature-sensitive for plants^45,46^ and Orchidaceae is usually high temperature-resistant, researchers could further determine the optimum temperature to improve the editing efficiency of DPII-Cpf1 system during orchid genetic transformation.

In summary, we successfully developed efficient multiplex genome editing tools (PTGm-Cas9 and DPII-Cpf1), and a protoplast-based screening system for *Phalaenopsis*. The protoplast-based screening platform provide a valuable foundation for developing more diverse and efficient genome editing toolkits for Orchidaceae, such as base editors and transcription regulation toolkits. Our study may also greatly promote the application of CRISPR/Cas multiplex genome editing technologies in Orchidaceae, facilitating large-scale orchid mutant library construction and orchid gene function and precise breeding studies.

## Materials and methods

### Plant materials and growth conditions

*Phalaenopsis* ST166 was purchased from Shenzhen Nongke Plant Clone Seedling Co., Ltd. The seedlings were cultured in illumination incubator at 25°C (light/dark photoperiod of 16 h/8 h).

### Protoplast isolation and transfection

The modified orchid protoplast isolation and transfection protocol was based on the method described by Yoo et al.^47^. Two-month-old *Phalaenopsis* ST166 seedling’s leaves were used for protoplast isolation. The leaves were cut into 0.5~1.0-mm strips using a fresh scalpel. The strips were transferred to a 60 mm petri dish containing 5 mL freshly prepared protoplast isolation solution (PIS). The PIS was made of 1% [w/v] Cellulase ‘Onozuka’ R–10 (Yakult Pharmaceutical), 0.2% [w/v] macerozyme R-10 (Yakult Pharmaceutical), 10 mM CaCl_2_ (Sigma, C5670), 0.4 M D-mannitol (Sigma, M1902), 20 mM KCl (Sigma, P5405), 0.1% BSA (Sigma, V900933), and 20 mM MES (pH 5.7, Sigma, M3671). The strips were digest for 3 h at 25 °C with gentle shaking in darkness. The protoplast suspension was then filtered through 40 μm nylon mesh to a 50 mL sterile tube, and wash the nylon mesh with equal volume of W5 solution, to remove tissue debris. The W5 solution contained 154 mM NaCl (Sigma, S5886), 5 mM KCl, 125 mM CaCl_2_, and 2 mM MES (pH 5.7). The solution was centrifuged at 100 g for 5 min at 22 °C, and removed the supernatant. The protoplast suspension was washed gently one more time with 5 mL W5 solution. Then the collected protoplasts were resuspended with 1 mL W5 solution. The protoplast cell concentration was measured using a hemocytometer. After counting, the protoplast suspension was centrifuged at 100 g for 3 min at 22°C, and resuspended with suitable volume of MMG solution to adjust the cell concentration about to 1×10^6^ cells/mL. The MMG solution contains 0.4 M D-mannitol, 5 mM MgCl_2_, and 4.0 mM MES, pH 5.7.

For protoplast transformation, the plasmid was delivered into phalaenopsis protoplast by PEG-mediated transfection method. 30 μg plasmid DNA (prepared by TIANGEN EndoFree Maxi Plasmid Kit, DP117) was used and mixed with 200 μL protoplast suspension. Then, equal volume (230 μL) of freshly prepared PEG solution was added into the mixture. The PEG solution contains 0.3 M D-mannitol, 100 mM CaCl_2_, and 30% PEG-4000 [w/v] (Sigma, 81240), PH 5.7. The transfection mixture was mixed gently and incubated at room temperature for 20 min. And then, the transfection reaction was stopped by adding 1 mL W5 solution. The mixture was centrifuged at 100 g for 3 min at 22°C to collect the protoplasts. The transfected protoplasts were gently resuspended with 1 mL W5 solution, and transferred to 6-well culture plate. After incubating 48 h at 25°C in darkness, protoplasts could be harvested for further experiments.

### Plasmid construction and extraction

For PTG-Cas9 and PTGm-Cas9 plasmids, the OsU3-PTG (Fig. S1) and OsU3-PTGm (Fig. S2) sequences were synthesized by BGI·Write, and the synthesized sequences were cut and inserted into the pYLCRISPR/Cas9Pubi-H binary vector^48^ using the two *Bsa* I sites.

For DPII-Cpf1 plasmid, first the *Phalaenopsis* codon-optimized LbCpf1 were synthesized by BGI·Write, and the LbCpf1 fragment was ligated into the pXZ binary vector, derived from pCAMBIA1300 vector, digested with *Hind* III/*EcoR* I. Second, the synthesized CmYLCV-HH-gRNAs-HDV (Fig. S3) was digested and inserted into the *Pst* I/*Xba* I sites of pXZ-Cpf1 vector, generated by the first step. Finally, the Super promoter was amplified with primers SuperP-F and SuperP-R (Table S1) and cloned into the plasmid pXZ-Cpf1-gRNAs generated by the previous step. The DPII-Cpf1 plasmid construction was finished and the multi-crRNAs could be replaced using the two *Aar* I sites.

### Protoplast direct PCR

The transformed protoplasts samples were centrifuged at 100 g for 3 min at 22°C to collect the protoplasts. And then the protoplasts were treated with 50 μL lysis buffer (20 mM Tris-HCl, 5 mM EDTA, 400 mM NaCl, 0.3% SDS, 200 μg/mL Proteinase K, PH 8.0) for DNA releasing at 55°C for 1 h. Following that, samples were treated at 95°C for 10 min. The lysed samples could be used for direct PCR amplification, using KOD FX Neo enzyme (TOYOBO, KFX-201). The PCR products were purified and used for library construction or other experiments.

### Mutation Detection and Analysis

Target sites of *PDS* gene were PCR-amplified using primers listed in Table S1. For high-throughput sequencing, the PCR products were purified with MinElute PCR Purification Kit (QIAGEN, 28006), and then were used for library construction using MGIEasy AmpSeq Library Prep Kit (MGI, 1000005257), and sequenced at BGISEQ-500 platform. Mutations were calculated based on the presence of mutations around the cleavage site. Specifically, the high quality clean data were obtained using fastp^49^, a robust FASTQ data pre-processing tool, to filter low quality reads, trim adapter and merge into a complete sequence. Bowtie 2^50^ was applied to align clean data to *PDS* gene sequence obtained by Sanger Sequencing. Mutation detection was analyzed using homemade well-packaged Python scripts.

## Supporting information

Supplemental meterials

## Author contributions

Xia K., Zhang D., and Liu G. performed the protoplast transformation and prepared the figures. Yang Y. performed the library construction. Xu X., and Zhang D. performed the sequencing data analysis. Gu Y., Sun H.-X., Zhang G. and Xia K. designed the experiments, interpreted the data and wrote the manuscript. All authors read and approved the final manuscript.

## Acknowledgments

This research was supported by China Postdoctoral Science Foundation (No. 2018M643197), and Guangdong Provincial Key Laboratory of Genome Read and Write (No. 2017B030301011), and Science, Technology and Innovation Commission of Shenzhen Municipality (No. JCYJ20170817151501595). We also sincerely thank the support provided by China National GeneBank (CNGB).

## Competing interests

The authors declare that they have no competing interests.

## Supplemental materials

Figure S1 The protoplast transient expression technology in *Phalaenopsis*.

Figure S2 PTGm-Cas9 system has a higher editing efficiency in *Phalaenopsis*.

Figure S3 PTG-Cas9 system with TaU6 promoter is effective in *Phalaenopsis*.

Figure S4 DNA sequence of multi-gRNA expression cassette of PTG-Cas9 system with OsU3 promoter.

Figure S5 DNA sequence of multi-gRNA expression cassette of PTG-Cas9 system with TaU6 promoter.

Figure DNA sequence of multi-gRNA expression cassette of PTGm-Cas9 system.

Figure S7 DNA sequence of multi-crRNA expression cassette of DPII-Cpf1 system.

Table S1. Primers used in this study.

## Data availability

The data that support the findings of this study have been deposited into CNGB Sequence Archive^51^ of CNGBdb^52^ with accession number CNP0001286.

## Notes

### Competing Interest Statement

The authors have declared no competing interest.

