## Supplemental meterials for "Efficient Multiplex Genome Editing Tools identified by Protoplast Technology in *Phalaenopsis*"

### Supplemental materials

Figure S1 The protoplast transient expression technology in *Phalaenopsis*.
Figure S2 PTGm-Cas9 system has a higher editing efficiency in *Phalaenopsis*.
Figure S3 PTG-Cas9 system with TaU6 promoter is effective in *Phalaenopsis*.
Figure S4 DNA sequence of multi-gRNA expression cassette of PTG-Cas9 system with OsU3 promoter.
Figure S5 DNA sequence of multi-gRNA expression cassette of PTG-Cas9 system with TaU6 promoter.
Figure DNA sequence of multi-gRNA expression cassette of PTGm-Cas9 system.
Figure S7 DNA sequence of multi-crRNA expression cassette of DPII-Cpf1 system.
Table S1. Primers used in this study.

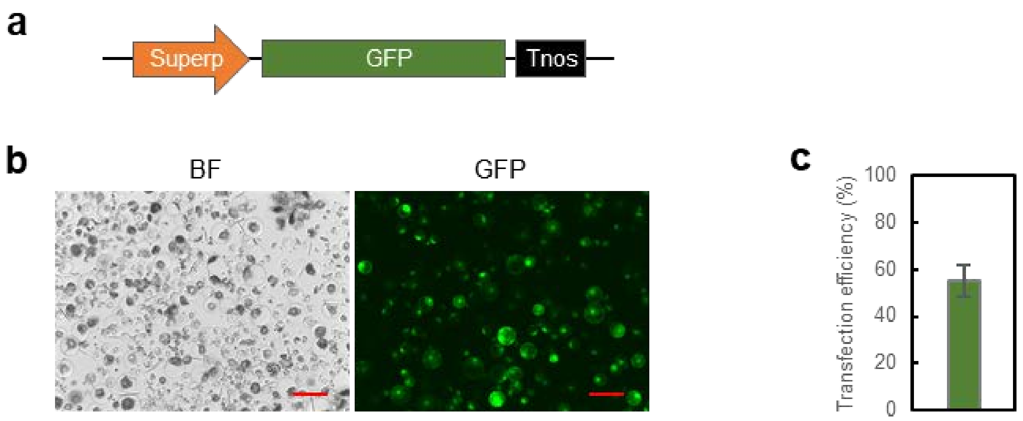


**Fig. S1 The protoplast transient expression technology in *Phalaenopsis*. a** The architecture of GFP expression cassette. **b** The transient expression of GFP protein in protoplasts of *Phalaenopsis* ST166. Bars = 200 μm. **c** The transfection efficiency of protoplast transient expression technology.

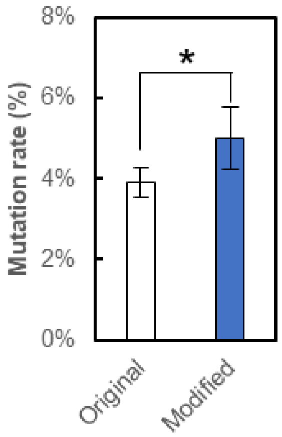


**Fig. S2 PTGm-Cas9 system has a higher editing efficiency in *Phalaenopsis*.** The sum of the mutation rates of the four sites. The Asterisk indicates a significant difference between original and modified sgRNA structure (Student's t test, *P<0.05).

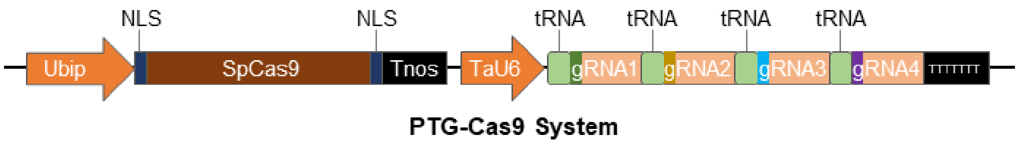


**b**

| Sample | Target ( mutation rate, %) | | | | |
|---|---|---|---|---|---|
| | 1 | 2 | 3 | 4 | 1~4 |
| T1 | 1.36 | 0.86 | 0.70 | 1.31 | 4.23 |
| T2 | 1.37 | 0.75 | 0.65 | 0.62 | 3.39 |

**Fig. S3 PTG-Cas9 system with TaU6 promoter is effective in *Phalaenopsis*. a** The architecture of PTG-Cas9 system with TaU6 promoter. **b** The mutation rate of the four target sites edited by PTG-Cas9 system with TaU6 promoter. The plasmid showed in a was delivered into *Phalaenopsis* protoplasts, and the editing results were analyzed by high-throughput sequencing. Mutation rate was calculated as the ratio of the number of mutant reads to that of the total reads.

**OsU3-PTG**

AGGAATCTTTAAACATACGAACAGATCACTTAAAGTTCTTCTGAAGCAACTTAAAGTTATCAGGC
ATGCATGGATCTTGGAGGAATCAGATGTGCAGTCAGGGACCATAGCACAAGACAGGCGTCTTC
TACTGGTGCTACCAGCAAATGCTGGAAGCCGGGAACACTGGGTACGTTGGAAACCACGTGTGA
TGTGAAGGAGTAAGATAAACTGTAGGAGAAAAGCATTTCGTAGTGGGCCATGAAGCCTTTCAG
GACATGTATTGCAGTATGGGCCGGCCCATTACGCAATTGGACGACAACAAAGACTAGTATTAGT
ACCACCTCGGCTATCCACATAGATCAAAGCTGGTTTAAAAGAGTTGTGCAGATGATCCGTGGCA
AACAAAGCACCAGTGGTCTAGTGGTAGAATAGTACCCTGCCACGGTACAGACCCGGGTTCGAT
TCCCGGCTGGTGCATCAAAGATGGCCTTTCTAGAGTTTTAGAGCTAGAAATAGCAAGTTAAAAT
AAGGCTAGTCCGTTATCAACTTGAAAAAGTGGCACCGAGTCGGTGCAACAAAGCACCAGTGGT
CTAGTGGTAGAATAGTACCCTGCCACGGTACAGACCCGGGTTCGATTCCCGGCTGGTGCAGGC
TTGCATAGCCTCTCAGGGTTTTAGAGCTAGAAATAGCAAGTTAAAATAAGGCTAGTCCGTTATCA
ACTTGAAAAAGTGGCACCGAGTCGGTGCAACAAAGCACCAGTGGTCTAGTGGTAGAATAGTAC
CCTGCCACGGTACAGACCCGGGTTCGATTCCCGGCTGGTGCAAAGCAATTTGTGCTGAGCAAGT
TTTAGAGCTAGAAATAGCAAGTTAAAATAAGGCTAGTCCGTTATCAACTTGAAAAAGTGGCACC
GAGTCGGTGCAACAAAGCACCAGTGGTCTAGTGGTAGAATAGTACCCTGCCACGGTACAGACC
CGGGTTCGATTCCCGGCTGGTGCAGGCAAATACATAAGCATCTCGTTTTAGAGCTAGAAATAGC
AAGTTAAAATAAGGCTAGTCCGTTATCAACTTGAAAAAGTGGCACCGAGTCGGTGCTTTTTTT

**Fig. S4 DNA sequence of multi-gRNA expression cassette of PTG-Cas9 system with OsU3 promoter.** OsU3 promoter and $tRNA^{Gly}$ is shown in orange and green respectively. The gRNA scaffold is highlighted with underline.

**TaU6-PTG**

GACCAAGCCCGTTATTCTGACAGTTCTGGTGCTCAACACATTTATATTTATCAAGGAGCACATTG
TTACTCACTGCTAGGAGGGAATCGAACTAGGAATATTGATCAGAGGAACTACGAGAGAGCTGA
AGATAACTGCCCTCTAGCTCTCACTGATCTGGGCGCATAGTGAGATGCAGCCCACGTGAGTTCA
GCAACGGTCTAGCGCTGGGCTTTTAGGCCCGCATGATCGGGCTTTGTCGGGTGGTCGACGTGTT
CACGATTGGGGAGAGCAACGCAGCAGTTCCTCTTAGTTTAGTCCCACCTCGCCTGTCCAGCAGA
GTTCTGACCGGTTTATAAACTCGCTTGCTGCATCAGACTTGAACAAAGCACCAGTGGTCTAGTG
GTAGAATAGTACCCTGCCACGGTACAGACCCGGGTTCGATTCCCGGCTGGTGCATCAAAGATG
GCCTTTCTAGAGTTTTAGAGCTAGAAATAGCAAGTTAAAATAAGGCTAGTCCGTTATCAACTTGA
AAAAGTGGCACCGAGTCGGTGCAACAAAGCACCAGTGGTCTAGTGGTAGAATAGTACCCTGCC
ACGGTACAGACCCGGGTTCGATTCCCGGCTGGTGCAGGCTTGCATAGCCTCTCAGGGTTTTAGA
GCTAGAAATAGCAAGTTAAAATAAGGCTAGTCCGTTATCAACTTGAAAAAGTGGCACCGAGTCG
GTGCAACAAAGCACCAGTGGTCTAGTGGTAGAATAGTACCCTGCCACGGTACAGACCCGGGTT
CGATTCCCGGCTGGTGCAAAGCAATTTGTGCTGAGCAAGTTTTAGAGCTAGAAATAGCAAGTTA
AAATAAGGCTAGTCCGTTATCAACTTGAAAAAGTGGCACCGAGTCGGTGCAACAAAGCACCAG
TGGTCTAGTGGTAGAATAGTACCCTGCCACGGTACAGACCCGGGTTCGATTCCCGGCTGGTGCA
GGCAAATACATAAGCATCTCGTTTTAGAGCTAGAAATAGCAAGTTAAAATAAGGCTAGTCCGTT
ATCAACTTGAAAAAGTGGCACCGAGTCGGTGCTTTTTTT

**Fig. S5 DNA sequence of multi-gRNA expression cassette of PTG-Cas9 system with TaU6 promoter.** TaU6 promoter and $tRNA^{Gly}$ is shown in orange and green respectively. The gRNA scaffold is highlighted with underline.

**OsU3-PTGm**

AGGAATCTTTAAACATACGAACAGATCACTTAAAGTTCTTCTGAAGCAACTTAAAGTTATCAGGC
ATGCATGGATCTTGGAGGAATCAGATGTGCAGTCAGGGACCATAGCACAAGACAGGCGTCTTC
TACTGGTGCTACCAGCAAATGCTGGAAGCCGGGAACACTGGGTACGTTGGAAACCACGTGTGA
TGTGAAGGAGTAAGATAAACTGTAGGAGAAAAGCATTTCGTAGTGGGCCATGAAGCCTTTCAG
GACATGTATTGCAGTATGGGCCGGCCCATTACGCAATTGGACGACAACAAAGACTAGTATTAGT
ACCACCTCGGCTATCCACATAGATCAAAGCTGGTTTAAAAGAGTTGTGCAGATGATCCGTGGCA
AACAAAGCACCAGTGGTCTAGTGGTAGAATAGTACCCTGCCACGGTACAGACCCGGGTTCGAT
TCCCGGCTGGTGCATCAAAGATGGCCTTTCTAGAGTTTCAGAGCTATGCTGGAAACAGCATAGC
AAGTTGAAATAAGGCTAGTCCGTTATCAACTTGAAAAAGTGGCACCGAGTCGGTGCAACAAAG
CACCAGTGGTCTAGTGGTAGAATAGTACCCTGCCACGGTACAGACCCGGGTTCGATTCCCGGCT
GGTGCAGGCTTGCATAGCCTCTCAGGGTTTCAGAGCTATGCTGGAAACAGCATAGCAAGTTGAA
ATAAGGCTAGTCCGTTATCAACTTGAAAAAGTGGCACCGAGTCGGTGCAACAAAGCACCAGTG
GTCTAGTGGTAGAATAGTACCCTGCCACGGTACAGACCCGGGTTCGATTCCCGGCTGGTGCAA
AGCAATTTGTGCTGAGCAAGTTTCAGAGCTATGCTGGAAACAGCATAGCAAGTTGAAATAAGGC
TAGTCCGTTATCAACTTGAAAAAGTGGCACCGAGTCGGTGCAACAAAGCACCAGTGGTCTAGTG
GTAGAATAGTACCCTGCCACGGTACAGACCCGGGTTCGATTCCCGGCTGGTGCAGGCAAATAC
ATAAGCATCTCGTTTCAGAGCTATGCTGGAAACAGCATAGCAAGTTGAAATAAGGCTAGTCCGT
TATCAACTTGAAAAAGTGGCACCGAGTCGGTGCTTTTTTT

**Fig. S6 DNA sequence of multi-gRNA expression cassette of PTGm-Cas9 system.** OsU3 promoter and $tRNA^{Gly}$ is shown in orange and green respectively. The modified gRNA scaffold is highlighted with an underline.

**CmYLCV-HH-crRNAs-HDV**

TGGCAGACATACTGTCCCACAAATGAAGATGGAATCTGTAAAAGAAAACGCGTGAAATAATGC
GTCTGACAAAGGTTAGGTCGGCTGCCTTTAATCAATACCAAAGTGGTCCCTACCACGATGGAAA
AACTGTGCAGTCGGTTTGGCTTTTTCTGACGAACAAATAAGATTCGTGGCCGACAGGTGGGGGT
CCACCATGTGAAGGCATCTTCAGACTCCAATAATGGAGCAATGACGTAAGGGCTTACGAAATAA
GTAAGGGTAGTTTGGGAAATGTCCACTCACCCGTCAGTCTATAAATACTTAGCCCCTCCCTCATT
GTTAAGGGAGCAAAATCTCAGAGAGATAGTCCTAGAGAGAGAAAGAGAGCAAGTAGCCTAGA
AGTAGTCAAGGCGGCGAAGTATTCAGGCACGTGGCCAGGAAGAAGAAAAGCCAAGACGACGA
AAACAGGTAAGAGCTAAGCTTCGCATGCCTCGTGAGACCGTCTTCGCAGGTGAAATTACTGATG
AGTCCGTGAGGACGAAACGAGTAAGCTCGTCTAATTTCTACTAAGTGTAGATATGGTAGCAGCA
ATAATGTCATTTAATTTCTACTAAGTGTAGATGTCAGGGAATAATGTTTCCAACTTAATTTCTACT
AAGTGTAGATCTTTACTCTGATCAGCAGCAATTTAATTTCTACTAAGTGTAGATAAGTATCATGTT
GTCAAAACACCGGCCGGCATGGTCCCAGCCTCCTCGCTGGCGCCGGCTGGGCAACATGCTTCG
GCATGGCGAATGGGACCACCTGCGAAGACGGTCTCGCGGTTCTAGAGTCGATCGACAAGTTTC
TCCATAATAATGTGTGAGTAGTTCCCAGATAAGGGAATTAGGGTTCCTATAGGGTTTCGCTCATG
TGTTGAGCATATAAGAAACCCTTAGTATGTATTTGTATTTGTAAAATACTTCTATCAATAAAATTT
CTAATTCCTAAAACCAAAATCCAGTACTAAAATCCAGATC

**Fig. S7 DNA sequence of multi-crRNA expression cassette of DP II -Cpf1 system.** CmYLCV promoter, HH and HDV are shown in orange, blue and green, respectively. The DR is highlighted with an underline.

**Table S1. Primers used in this study**

| Primer used for plasmid construction | |
|---|---|
| Name | Primer sequence (5’→3’) |
| SuperP-F | GCCAGTGCCAAGCTTGAGCTCACGGTATCGATAAGCTC |
| SuperP-R | TCTTCTTTGGAGCCATGGTACCCTAGAGTCGATTTGGT |
| Primers used for target region amplification | |
| Name | Primer sequence (5’→3’) |
| F1 | ATCCAAATTATCAATTACCCA |
| R1 | TCAATCTTTTGTACACGAGAA |
| F2 | CACATTGAGTCACTAGGCGGACA |
| R2 | ATTCAAATTCACAGAACAATTTC |
| F3 | CTTCGCGCCAGCAGAAGAAT |
| R3 | CGAACATATTTAATTTATGATTTTAGAACTCACCTT |
